# The sustained release of basic fibroblast growth factor accelerates angiogenesis and the survival of the inactivated dermis by high hydrostatic pressure

**DOI:** 10.1101/477042

**Authors:** Tien Minh Le, Naoki Morimoto, Toshihito Mitsui, Sharon Claudia Notodihardjo, Maria Chiara Munisso, Natsuko Kakudo, Kenji Kusumoto

**Affiliations:** Department of Plastic and Reconstructive Surgery, Kansai Medical University, Hirakata, Osaka, Japan

## Abstract

We developed a novel skin regeneration therapy combining nevus tissue inactivated by high hydrostatic pressure (HHP) in the reconstruction of the dermis with a cultured epidermal autograft (CEA). The issue with this treatment is the unstable survival of CEA on the inactivated dermis. In this study, we applied collagen/gelatin sponge (CGS), which can sustain the release of basic fibroblast growth factor (bFGF), to the inactivated skin in order to accelerate angiogenesis. Murine skin grafts from C57BL6J/Jcl mice (8 mm in diameter) were prepared, inactivated by HHP and cryopreserved. One month later, the grafts were transplanted subcutaneously onto the back of other mice and covered by CGS impregnated with saline or bFGF. Grafts were taken after one, two and eight weeks, at which point the survival was evaluated through the histology and angiogenesis-related gene expressions were determined by real-time polymerase chain reaction. Histological sections showed that the dermal cellular density and newly formed capillaries in the bFGF group were significantly higher than in the control group. The relative expression of FGF-2, PDGF-A and VEGF-A genes in the bFGF group was significantly higher than in the control group at Week 1. This study suggested that the angiogenesis into grafts was accelerated, which might improve the survival in combination with the sustained release of bFGF by CGSs.

## Introduction

Basic fibroblast growth factor (bFGF) is an essential mitogen that plays a crucial role in the wound healing processes by not only stimulating cell growth and differentiation but also inducing neovascularization and connective tissue synthesis [1–3]. As human recombinant bFGF has been commercially available in Japan since 2001, its topical administration has been shown to be effective for wound healing in clinical treatments and has recently been applied more extensively [4–6]. However, the disadvantage of this medication is the need for its daily administration on the wound due to its short half-life *in vivo* [2]. To overcome this issue, we developed a novel collagen/gelatin scaffold (CGS) containing 10wt% acidic gelatin that is capable of the sustained release of a charged growth factor, such as bFGF [7], platelet-derived growth factor (PDGF) [8] or hepatocyte growth factor (HGF) [9] for more than 10 days. We previously showed that CGS impregnated with bFGF at 7-14 μg/cm^2^ achieved the most efficient promotion of angiogenesis and dermis-like tissue formation, even in a delayed wound-healing model of diabetic mice [10–12].

Recently, a number of different decellularization methods, including chemical, biological and physical and miscellaneous agents, have been suggested to be useful for generating superior bioengineering tissue [13]. Decellularized tissue retains its native structure or mechanical properties and has been reported to show low or no immunogenicity [14, 15], making it an ideal substitute or scaffold for tissue engineering and regenerative medicine. High hydrostatic pressure (HHP), another potential new decellularization method, is a physical technique that can inactivate cells or tissues in a short time without using any chemical reagents [16–20]. We reported that HHP treatment exceeding 200 MPa for 10 min was sufficient to induce cell death through the inactivation of mitochondrial activity, resulting in the complete inactivation of human skin, human nevus specimens and porcine skin without damaging the extracellular matrix (ECM) [21–23].

Based on these principles, we developed a novel treatment for giant congenital melanocytic nevi (GCMN) that involves the reuse of inactivated autologous nevus in combination with cultured epidermal autograft (CEA) using Green’s method [24–26]. However, the infiltration of fibroblasts into the inactivated skin took more than a week [27], and the thickness of the inactivated skin decreased at 12 weeks after implantation [23, 24].

Therefore, in the present study, we investigated the promotion of angiogenesis at inactivated dermis by applying CGS impregnated with bFGF. The murine skin was inactivated by HHP at 200 MPa and implanted subcutaneously in combination with a bFGF-impregnated CGS. The angiogenesis-related gene expression and the survival of the inactivated dermis were then evaluated after the implantation.

## Materials and methods

### Ethics statement

All animal experiments in this study were conducted at Kansai Medical University in accordance with the Guidelines for Animal Experiments established by the Ministry of Health, Labor and Welfare of Japan and by Kansai Medical University, Japan. The number of animals used in this study was kept to a minimum, and the protocol was approved by the Animal Care and Use Committee of Kansai Medical University (permit No. 17-002(02)). The experiment protocol was also approved by the Ethics Committee of Kansai Medical University (permit No. 1472).

### Preparation of inactivated murine skin grafts

Ten 8-week-old male C57BL6J/Jcl mice (CLEA Japan Inc., Tokyo, Japan) were euthanized by carbon dioxide inhalation, after which the fur on their backs was shaved with an electric razor (Thrive; Daito Electric Machine Ind. Co., Ltd., Japan), and the skin was depilated using hair-removing body foam (Kracie, Tokyo, Japan). Next, the whole back skin was collected, and rounded 8-mm full-thickness skin grafts were obtained using punch biopsy tools (Kai Industries Ltd., Gifu, Japan). Twenty grafts were packed into each of six 25-ml cryopreservation bottles (Perfluoroalkoxy [PFA] bottle; Tech-jam, Osaka, Japan) filled with 10% glycerol + 5% fructose solution (GLYCEOL^®^; Chugai Pharm, Ltd., Tokyo, Japan) (total of 120 skin grafts).

The skin grafts in each bottle were pressurized at 200 MPa for 10 minutes using an custom made HHP device (Echigo Seika Co., Ltd., Nagaoka, Japan), as described in our previous studies [22, 26, 28]. This system consists of an isostatic chamber, a main pressure unit with a hydraulic hand pump, an electric pressure generation unit and a pressure control unit (Fig 1). Finally, skin specimens were cryopreserved in freezer at −80 °C until grafting.

**Fig 1:**
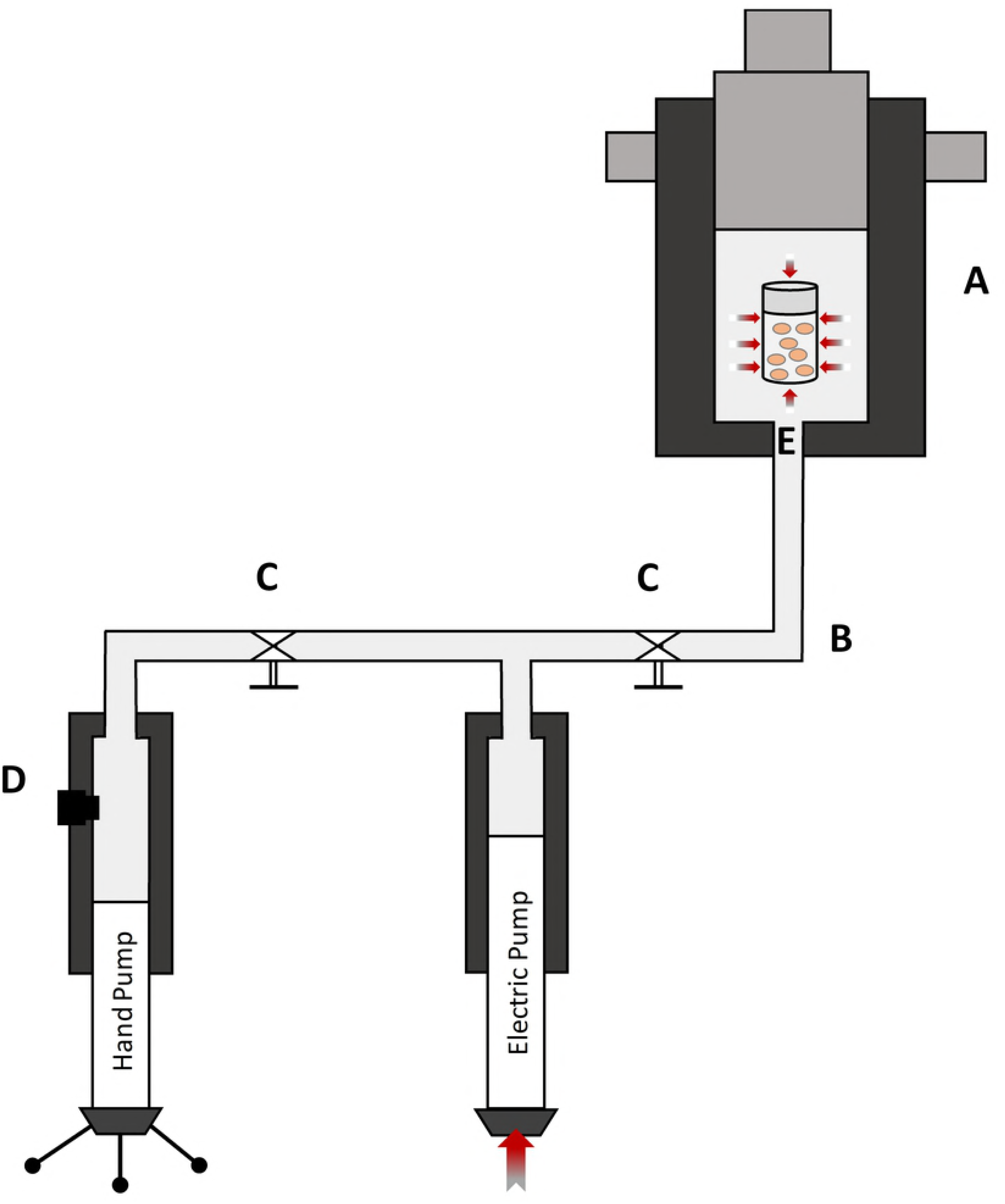
Schematic representation of HHP device activity. (A) isostatic chamber, (B) liquid pipelines, (C) valves, (D) trap-air releasing, (E) pressure and temperature sensors.

To assess the effect of HHP on murine skin, the full-thickness grafts before and after HHP treatment as well as four weeks after cryopreservation were subjected to hematoxylin-eosin (HE) and Azan staining (Fig 2A, 2B, 2C).

**Fig 2.**
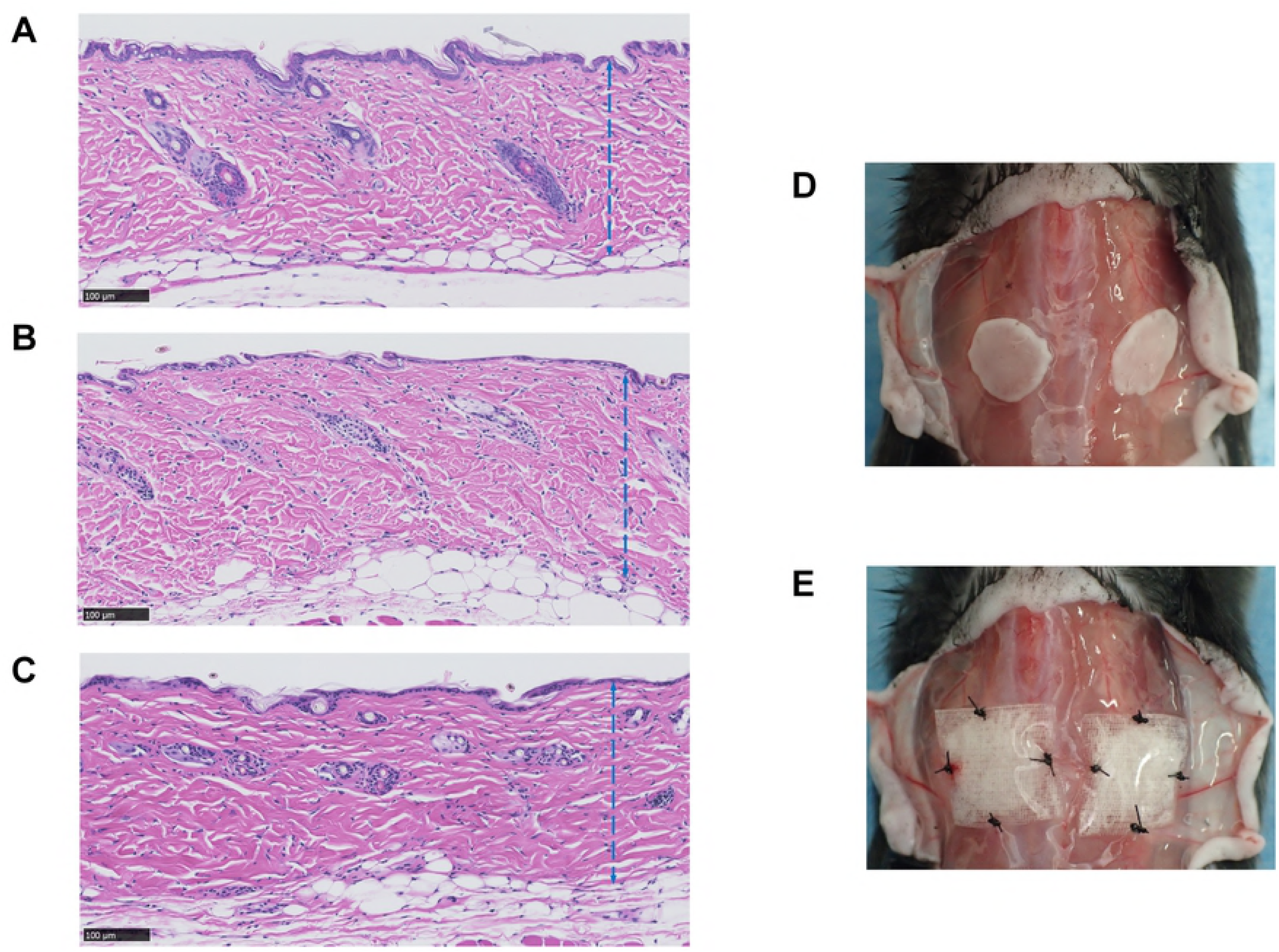
Micrographs of the inactivated skin grafts by HHP and photographs of grafting procedures. (A) A micrograph of murine skin before HHP, (B) a micrograph of murine skin just after HHP, (C) a micrograph of murine skin after cryopreservation for 4 weeks; the blue broken lines indicate the dermis of grafts, (D) a gross photo of the inactivated graft position, (E) gross photos of the grafts covered by NSS- or bFGF-impregnated (sandwich-technique) CGSs. Magnification: 20x, scale bar: 100 μm.

### Impregnation of CGS with NSS and bFGF

The CGSs used in this study were kindly supplied by Gunze Co. Ltd. (Ayabe, Japan) [7, 11]. CGSs 1 × 1 cm in size were placed on 10-cm tissue culture dishes (Falcon; Corning Inc., Corning, NY, USA). Human recombinant bFGF (500 μg; Fiblast Spray^®^; Kaken Pharmaceutical, Tokyo, Japan) was dissolved in 7.14 ml of normal saline solution (NSS; Fuso Pharmaceutical Industries, Ltd., Osaka, Japan), and 70 μg/ml of bFGF solution was prepared. Each CGS in the NSS group was injected with 100 μl of NSS using a micropipette, while in the bFGF group, each CGS was injected with 100 μl of the prepared bFGF solution to achieve a bFGF concentration of 7 μg/cm^2^. The sponges were then incubated for 30 min at room temperature for impregnation, after which they were stored in refrigerator at 4 °C overnight until implantation. Fifty CGSs each impregnated with either NSS or bFGF were prepared for grafting procedures.

### Implantation of inactivated skin grafts in combination with CGSs

We used totally 40 eight-week-old male mice (C57BL/6JJcl, CLEA Japan Inc., Tokyo, Japan) for the grafting procedure. Anesthesia was performed using 5% isoflurane inhalation (Wako Pure Chemical Industries Ltd., Osaka, Japan) delivered by a vaporizer (Forawick Vaporizer; Muraco Medical Co., Ltd., Tokyo, Japan) at a standardized concentration through the outlet tube. For induction, animals were placed in a glass box (22 × 9 × 11 cm) connected by a silicon tube to an anesthetic gas mixture of 3% vaporized isoflurane for 60 seconds. For maintenance, 1.5% to 2% isoflurane was used during the surgical procedures. After the back skin had been shaved and depilated, it was incised longitudinally, and the deep fascia of the back region was exposed. Cryopreserved inactivated skin grafts were thawed in a water bath at 37 °C, and 2 grafts per mouse were grafted onto the dorsal fascia symmetrically and sutured with 5-0 nylon sutures (Bear Corporation, Osaka, Japan). Subsequently, in the bFGF group (n=20, 40 grafts), a CGS impregnated with bFGF was placed onto each graft in a sandwich technique and sutured to the fascia using nylon 5-0 thread to ensure the sustained release of the growth factor (Fig 2D, 2E). Finally, the skin was closed with nylon 5-0 sutures. Twenty mice in the bFGF group were ultimately treated in this manner. In the NSS group (n=20, 40 grafts), CGSs impregnated with NSS were placed onto the skin grafts via the same procedure.

### Evaluating angiogenesis related genes expression of cells infiltrating the grafts

One and two weeks after implantation, four mice in each group were euthanized via carbon dioxide inhalation. Skin grafts (n=2 per mouse, n=8 per group) were taken, and the connective tissue were removed. The grafts were instantly frozen in liquid nitrogen to prevent RNA degradation. Each frozen tissue sample was pulverized to a fine powder with a standard liquid nitrogen pre-chilled mortar and pestle. This powder was then transferred to an RNase-free 1.5-ml tube (Eppendorf^®^ DNA LoBind tubes; Eppendorf Co., Hamburg, Germany) and homogenized with 350 μl lysate solution, after which the total RNA was purified by column precipitation using a NucleoSpin^®^ RNA Plus (Macherey-Nagel GmbH & Co. KG, Düren, Germany) according to the manufacturer’s instructions. The extracted total RNA was quantitated by measuring the absorbance in a nanodrop spectrophotometer (Nanodrop ND-1000; Thermo Fisher Scientific, Wilmington, De, USA). The absorbance ratio at A260/A280 nm of all samples ranged from 1.8 to 2.1, indicating that they were pure during the RNA extraction procedures. The quality and purity of the extracted RNA were also determined by electrophoresis on a 1.5% (w/v) agarose gel containing ethidium bromide (EtBr) and visualized under UV light (FAS-IV; Nippon Genetics Co., Ltd., Tokyo, Japan).

RT-PCR was performed to detect the expression of growth factors related to angiogenesis, including FGF-2, PDGF-A and vascular endothelial growth factor-A (VEGF-A). Glyceraldehyde-3-phosphate dehydrogenase (GAPDH) was used as reference gene [29], and a total of four primers (Table 1) were obtained in a commercially available Quantitect^®^ Primer Assay (Qiagen Inc., Hilden, Germany). A total of 25 μl of reaction sample, comprising 10 ng of total RNA samples, primers, Rotor-Gene SYBR Green RT-PCR Master Mix, and Rotor-Gene RT Mix (Rotor-Gene SYBR^®^ Green RT-PCR Kit; Qiagen Inc.), was loaded onto a 72-well Rotor Disc (Qiagen Inc.). The amplification protocol was run using a Real-Time One Step RT-PCR with Rotor-Gene Q device (Qiagen Inc.) according to the manufacturer’s conditions: 55 °C/10 min, 95 °C/5 min, 40 cycles at 95 °C/5 sec, 60 °C/10 sec and melting. The specificity and PCR efficiency of each primer was confirmed by a single-peak melting curve analysis and standard curve of 5-fold serial dilutions, and the results were quantified using the comparative 2^-ΔΔCt^ method [30, 31]. Each individual sample was tested in duplicate, and the expression of the grafts in the NSS group at each time point was normalized to 1.

**Table 1.** Primer details

| Gene name | Gene symbol | Catalogue number | Accession number | Product length |
| --- | --- | --- | --- | --- |
| Fibroblast growth factor 2 | Fgf2 | QT00128135 | NM_008006 | 138 |
| Platelet derived growth factor A | Pdgfa | QT00197610 | NM_008808 | 108 |
| Vascular endothelial growth factor A | Vegfa | QT00160769 | NM_009505 | 117 |
| Glyceraldehyde-3-phosphate dehydrogenase | Gapdh | QT01658692 | NM_008084 | 144 |

### Sampling and histological evaluation of the skin grafts

At one, two and eight weeks after implantation, four mice per group were euthanized using carbon dioxide inhalation. The implanted grafts (n=8 per group) including the surrounding tissue (2 × 2 cm) and muscle underneath were harvested. Tissue specimens were cut axially at the center using a scalpel, fixed with 10% formalin-buffered solution (Wako Pure Chemical Industries, Ltd.), and then embedded in paraffin blocks. Paraffin sections 5-μm-thick from the central region of each specimen were subjected to HE staining, Azan staining and immunohistochemical staining for anti-vimentin and anti-CD31. After staining, histological photographs were taken and analyzed using a NanoZoomer 2.0 HT whole-slide imager with the NDP.view2 software program (Hamamatsu Photonics, Hamamatsu, Japan) at 40× magnification.

### The histological evaluation of the cellular density of grafts

The number of cells that had infiltrated the grafts was evaluated on HE sections at Weeks 1 and 2. The number of cell nuclei in 3 areas measuring 250 × 250 μm (at the center and on both sides 500 μm from the center) were counted in each section and compared.

### The immunohistochemical evaluation of the newly formed capillaries and infiltrating fibroblasts in the grafts

Immunohistochemical staining of anti-vimentin was performed to detect infiltrated fibroblasts, and anti-CD31 immunohistochemistry was used to evaluate newly formed capillaries in the grafts at Weeks 1 and 2. For staining, 5-μm-thick sections were deparaffinized and rehydrated, and then antigen retrieval processing was performed using the heat-induced target retrieval method. Sections were immersed in a pre-heated staining dish containing EDTA buffer (pH 9; Nichirei Biosciences Inc., Tokyo, Japan) and incubated at 98 °C for 20 min (anti-CD31) or in a dish containing 10 mM sodium citrate buffer (pH 6.0; Sigma-Aldrich Japan. Co. Ltd., Tokyo, Japan) and incubated at 98 °C for 40 min (anti-vimentin). After being cooled to room temperature, the sections were rinsed twice in distilled water (DW) and immersed in 3% hydrogen peroxide (H_2_O_2_; Wako Pure Chemical Industries, Ltd.) for 10 min to block endogenous peroxidase activity. After rinsing with DW and TBST (50 mM Tris-HCl buffer, pH 7.6, containing 0.05% tween-20 and 0.15 M NaCl; Nichirei Biosciences Inc.), the protein blocking solution (3% bovine serum albumin in PBS; Sigma-Aldrich Japan Co., Ltd.) was applied to sections for 60 min to block non-specific protein binding. Subsequently, the sections were incubated with rabbit polyclonal anti-CD31 antibody (dilution 1:300; Spring Bioscience, Fremont, CA, USA) or rabbit monoclonal anti-vimentin antibody (dilution 1:1000, Abcam Co., Ltd., Boston, MA, USA) as a primary antibody and incubated at 4 °C overnight. Sections were then rinsed in TBST, and the peroxidase-labeled secondary antibody (Histofine^®^ Simple Stain MAX PO^®^; Nichirei Biosciences Inc.) was applied at room temperature for 60 min (anti-CD31) or 30 min (anti-vimentin). Subsequently, the sections were rinsed in TBST again before being exposed to 3,3’-diaminobenzidine tetrahydrochloride (DAB; Nichirei Biosciences Inc.) and counterstained with hematoxylin.

In anti-CD31 immunohistochemical sections, the numbers and area of the newly formed capillaries in the dermal part with a clearly visible lumen and erythrocytes inside were counted manually using the NDP.view2 software program (Hamamatsu Photonics), and the sum was used for the data analysis. In anti-vimentin immunohistochemical sections, typically flattened or extensible shaped fibroblasts in 3 areas measuring 250 × 250 μm (at the center and on both sides 500 μm from the center) were counted in each section, and the dermal fibroblast density was calculated and compared.

### The histological evaluation of the graft thickness

The thickness of each graft after 8 weeks was evaluated on Azan-stained sections. Azan staining stains dermal collagen fibers a dark blue that can be differentiated from the connective tissue stained a light blue. The maximal dermal thickness of the right third, central third, and left third as well as that of each graft were estimated for these sections and compared. The cross-sectional area of the dermis also measured on Azan-stained sections at Week 8 using the NanoZoomer 2.0 HT whole-slide imager with the NDP.view2 software program (Hamamatsu Photonics).

### Statistical analysis

Statistical analyses were performed using the Microsoft Excel (Office 365; Microsoft Corporation, Redmond, WA, USA) and Prism 7.03 (GraphPad Software, Inc., San Diego, CA, USA) software programs. A one-way analysis of variance (ANOVA), followed by Tukey-Kramer post hoc test, was used to compare cells infiltration and angiogenesis for each condition and experimental time point; Student’s *t*-test was used when comparing two groups. All data are presented as the mean ± standard error of mean (SEM), and a p-value < 0.05 was considered statistically significant.

## Results

### The histological evaluation of the grafts

The micrographs of HE staining sections of the skin grafts before implantation are shown in Fig 2A, 2B, 2C. No marked differences in the epidermis layer, dermal collagen fibers, cell nuclei or other adnexal structures were noted after pressurization or cryopreservation. At Week 1, the epidermis was detached from the dermis (Fig 3A), and the original graft’s cell remnants were absorbed. Recipient cells then infiltrated the grafts from the fascia, and the epidermis was completely removed; no severe inflammation response or degradation of dermal collagen fibers was observed at Week 2. A comparison of the dermal cellular density (Fig 3B) showed that the number of cells in the bFGF group significantly increased from Weeks 1 to 2 (p < 0.001), and significant differences were noted in the density between the bFGF and NSS groups at Week 2 (p < 0.01). Anti-vimentin staining was used to differentiate the infiltrated fibroblasts from other cells in the grafts (Fig 4A). The micrographs of anti-vimentin-stained sections show that many fibroblasts had already migrated into the grafts at Week 1. The number of infiltrated fibroblasts increased at Week 2 in both groups, and the fibroblast density in the bFGF group at Week 2 was significantly higher than that in the NSS group (p < 0.001) (Fig 4B).

**Fig 3.**
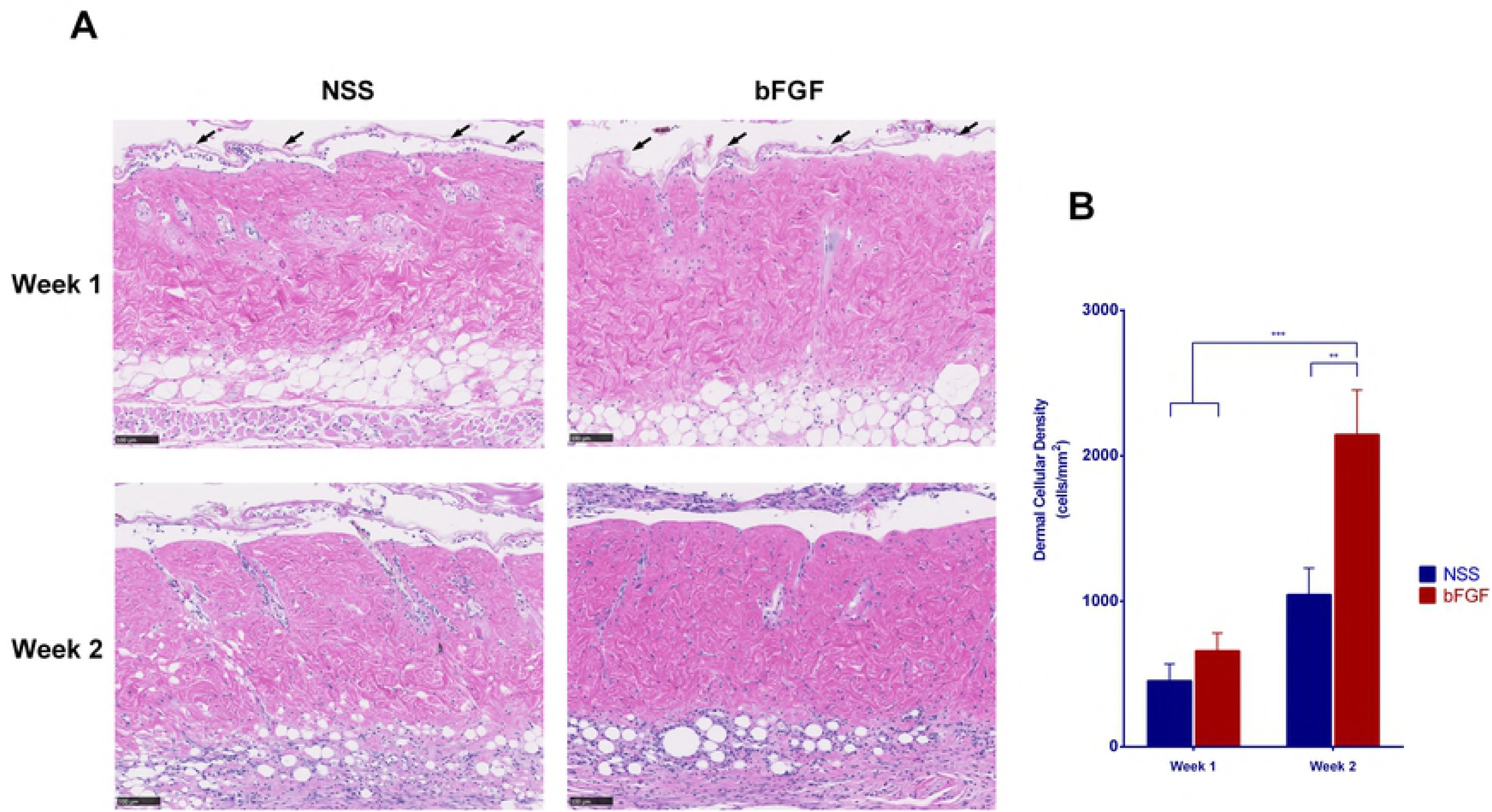
The infiltration of recipient cells into inactivated dermis after implantation. (A) Micrographs of HE-stained sections of skin grafts at Weeks 1 and 2 after implantation. The original dermal cell remnants were absorbed, and the epidermis indicated by black arrowheads was detached from the papillary dermis at Week 1. Cells from recipients showed infiltration to the dermis, and the dermal cellular density increased over the time. No inflammation reactions or dermal collagen fiber degeneration was observed. Magnification: 10x, scale bar: 100 μm. (B) The diagram shows significant differences in the dermal cellular density of the bFGF group between Weeks 1 and 2 (*** p < 0.001); the dermal cellular density of the bFGF group at Week 2 was significantly higher than that in the NSS group (** p < 0.01).

**Fig 4.**
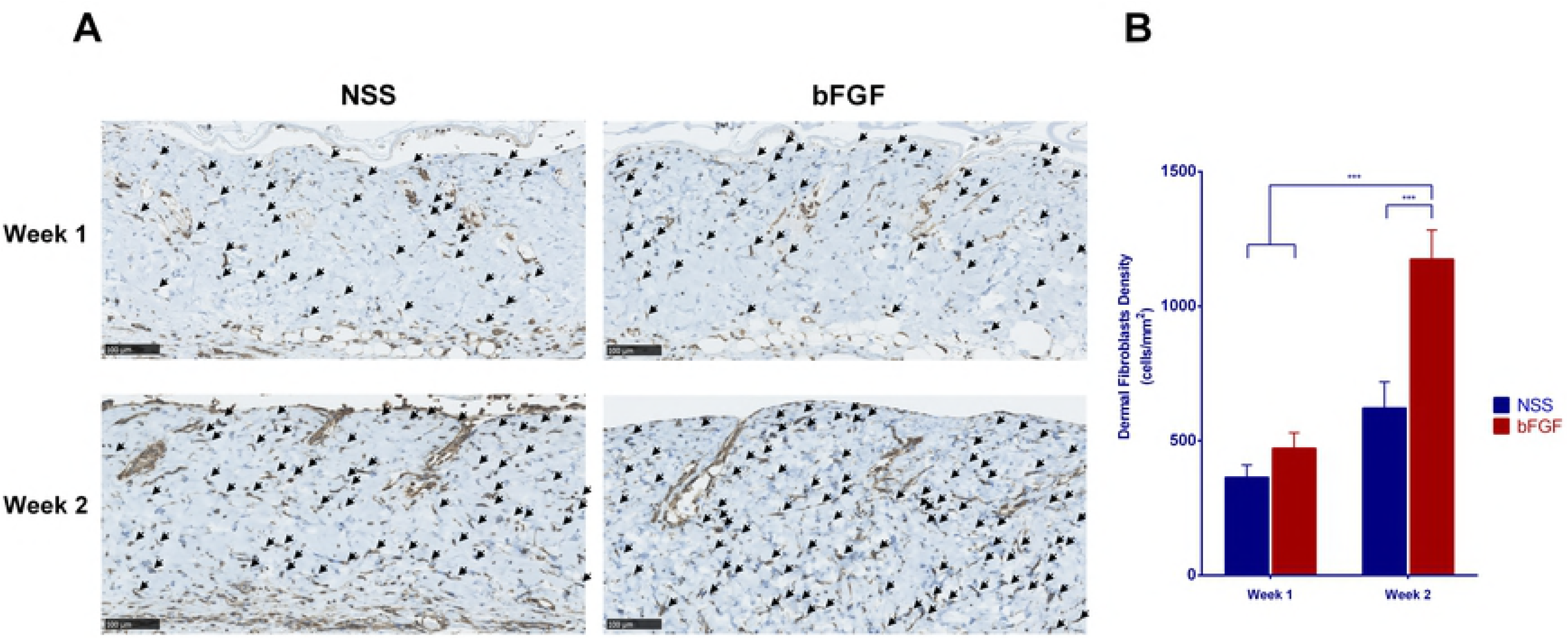
The infiltration of recipient fibroblasts into inactivated dermis after implantation. (A) Micrographs of anti-vimentin-stained sections differentiating fibroblasts from other cells at Weeks 1 and 2 after implantation. Recipient fibroblasts began to infiltrate the grafts at Week 1 and increased at Week 2. ↙: fibroblast; magnification: 20x, scale bar: 100 μm. (B) A comparison of the dermal fibroblast density of the grafts; a significant increase in the number of fibroblasts in the bFGF group was noted between Weeks 1 and 2 as well as between the bFGF and NSS groups in Week 2 (*** p < 0.001).

### The assessment of RT-PCR for FGF-2, PDGF-A and VEGF-A

To evaluate the effects of bFGF impregnation of CGS on dermal angiogenesis, the mRNA expression of endogenous FGF-2, PDGF-A and VEGF-A were measured by real-time PCR. At Week 1, the expression of endogenous FGF-2 in the dermis of the bFGF group was elevated and significantly higher (17-fold) than that in the NSS group (p < 0.001). The expression of PDGF-A and VEGF-A were also increased in the bFGF group (38- and 44-fold, respectively) compared with the NSS group (p < 0.01, Fig 5A). However, after 2 weeks, the levels of FGF-2 in the bFGF group were significantly down-regulated compared with the NSS group (p < 0.01), and the levels of PDGF-A in the bFGF group were also reduced at that point, although they remained 2-fold higher than in the NSS group (p < 0.05, Fig 5B).

**Fig 5.**
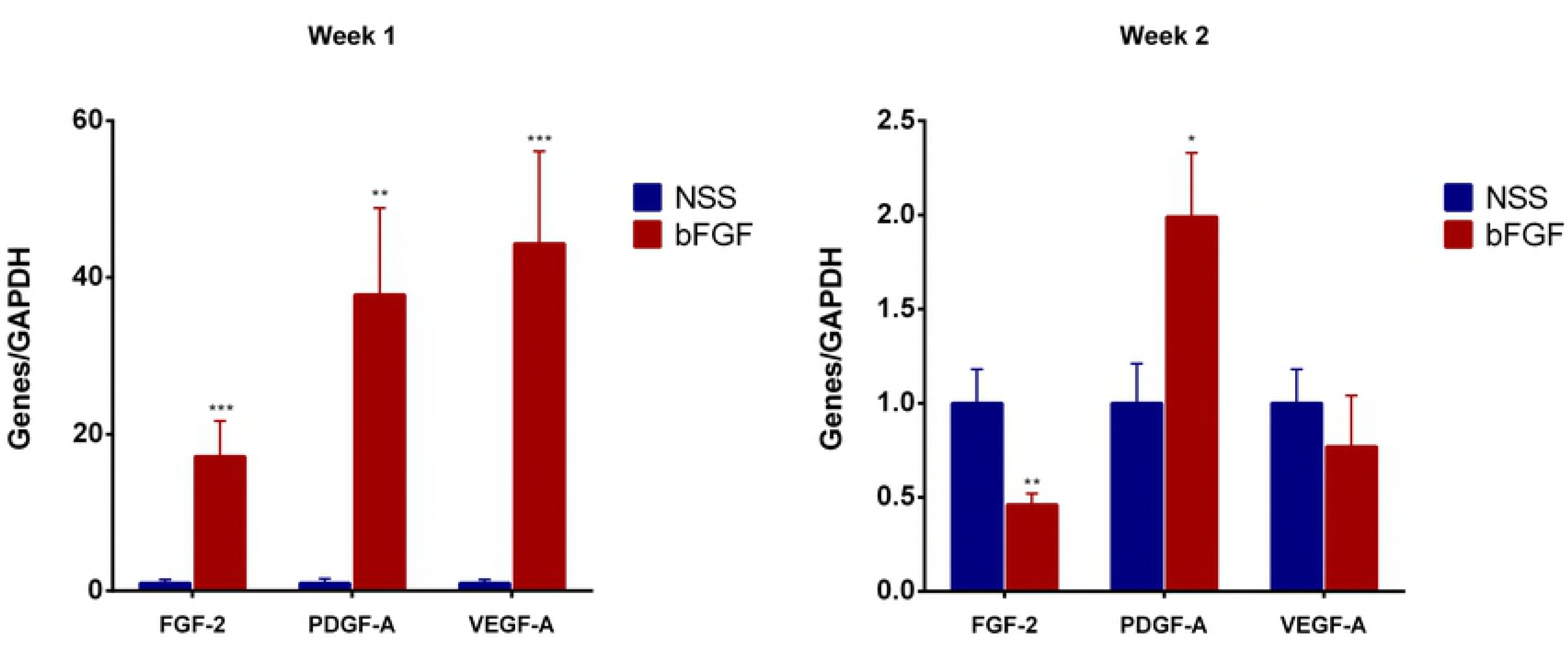
The comparison of the angiogenesis-related genes expression at Weeks 1 and 2 after implantation. The relative expressions of FGF-2, PDGF-A and VEGF-A were strongly up-regulated in the bFGF group at Week 1 (** p < 0.01, *** p < 0.001). The expression of FGF-2 was lower in the bFGF group than in the NSS group (** p < 0.01), while that of PDGF-A was higher in the bFGF group than in the NSS group (* p < 0.05).

### The immunohistological evaluation of the newly formed capillaries

Micrographs of the anti-CD31 immunohistochemical staining of grafts at Weeks 1 and 2 are shown in Fig 6A. This staining detected capillaries (black arrowhead) in the grafts of both groups, revealing the blood flow to the grafts. Only a few small capillaries were observed at Week 1 in both groups. However, the capillary formation increased in number and size at Week 2, especially in the bFGF group. The comparison of the number of newly formed capillaries in the grafts (Fig 6B) showed that the number of capillaries increased over time in both the bFGF group (p < 0.01) and NSS group (p < 0.05). The number of capillaries in the bFGF group was significantly higher than in the NSS group at Weeks 1 (p < 0.01) and 2 (p < 0.05). The comparison of the area of newly formed capillaries in the grafts (Fig 6C) also showed that the number of capillaries increased over time in both the bFGF group (p < 0.05) and NSS group (p < 0.05). A significant difference in this number was observed between the groups (p < 0.05) at Weeks 1 and 2.

**Fig 6.**
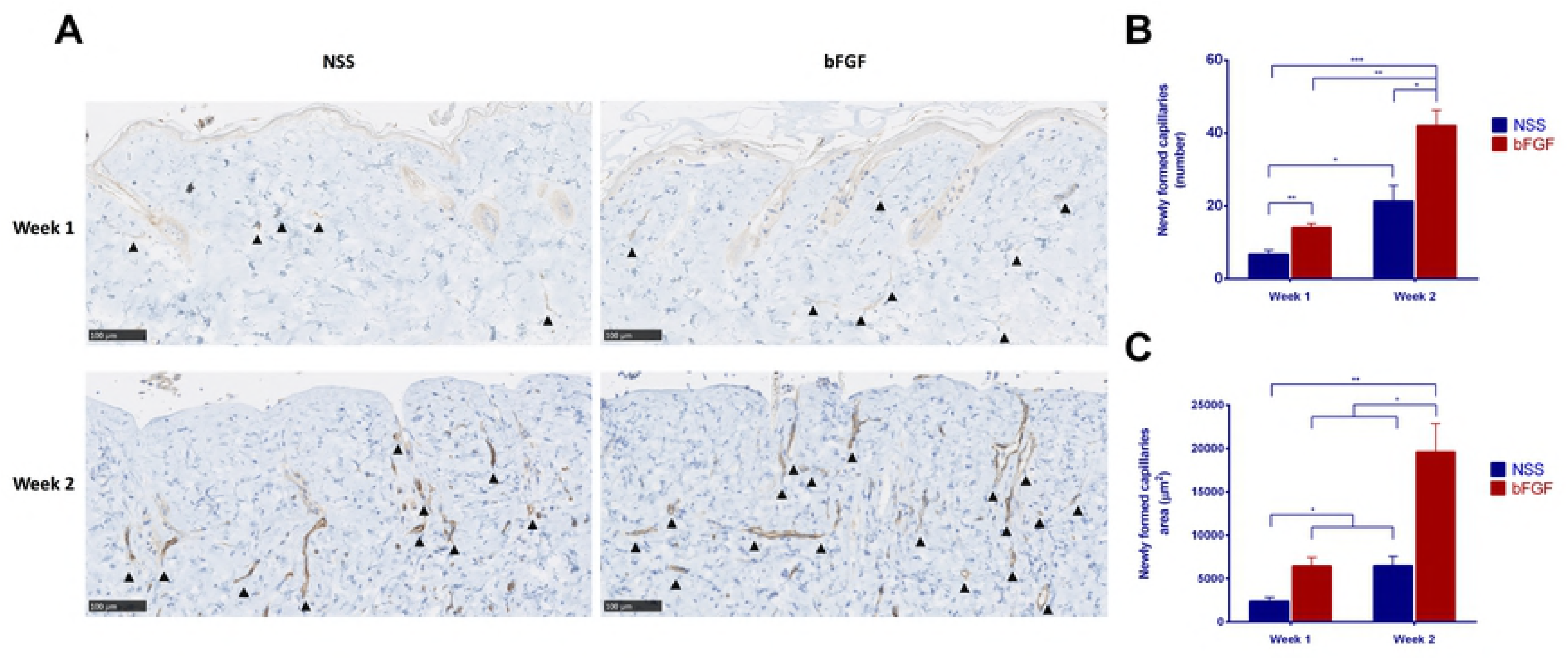
Evaluation of newly formed capillaries after implantation. (A) Micrographs of immunohistochemical staining of anti-CD31 at Weeks 1 and 2. Few capillaries were observed at Week 1 in both groups. The capillaries increased in number and size at Week 2 in both groups. ▲: newly formed capillary; magnification: 20x, scale bar: 100 μm. (B) A comparison of the number of newly formed capillaries in the grafts. The number of capillaries significantly increased from Weeks 1 to 2 in both groups. The number of capillaries in the bFGF group was significantly higher than that in the NSS group at Weeks 1 and 2 (* p < 0.05, ** p < 0.01, *** p < 0.001). (C) A comparison of the newly formed capillaries area in the grafts. The area of the capillaries also significantly increased from Weeks 1 to 2 in both groups. The number of capillaries in the bFGF group was significantly higher than that in the NSS group at Weeks 1 and 2 (* p < 0.05, ** p < 0.01).

### The assessment of the thickness and area of the grafts

Micrographs of Azan-stained section of skin grafts at eight weeks after implantation are shown in Fig 7A. Whole cross-sections of each graft are shown, with the dermal part stained dark blue and indicated by a broken black line. The thickness in the bFGF group seemed greater than that in the NSS group, and the dermal area also seemed to be larger in the bFGF group. Comparing the thickness and area of the grafts at Week 8 showed that the thickness and area were significantly greater (Fig 7B, 7C) in the bFGF group than in the NSS group (p < 0.05).

**Fig 7.**
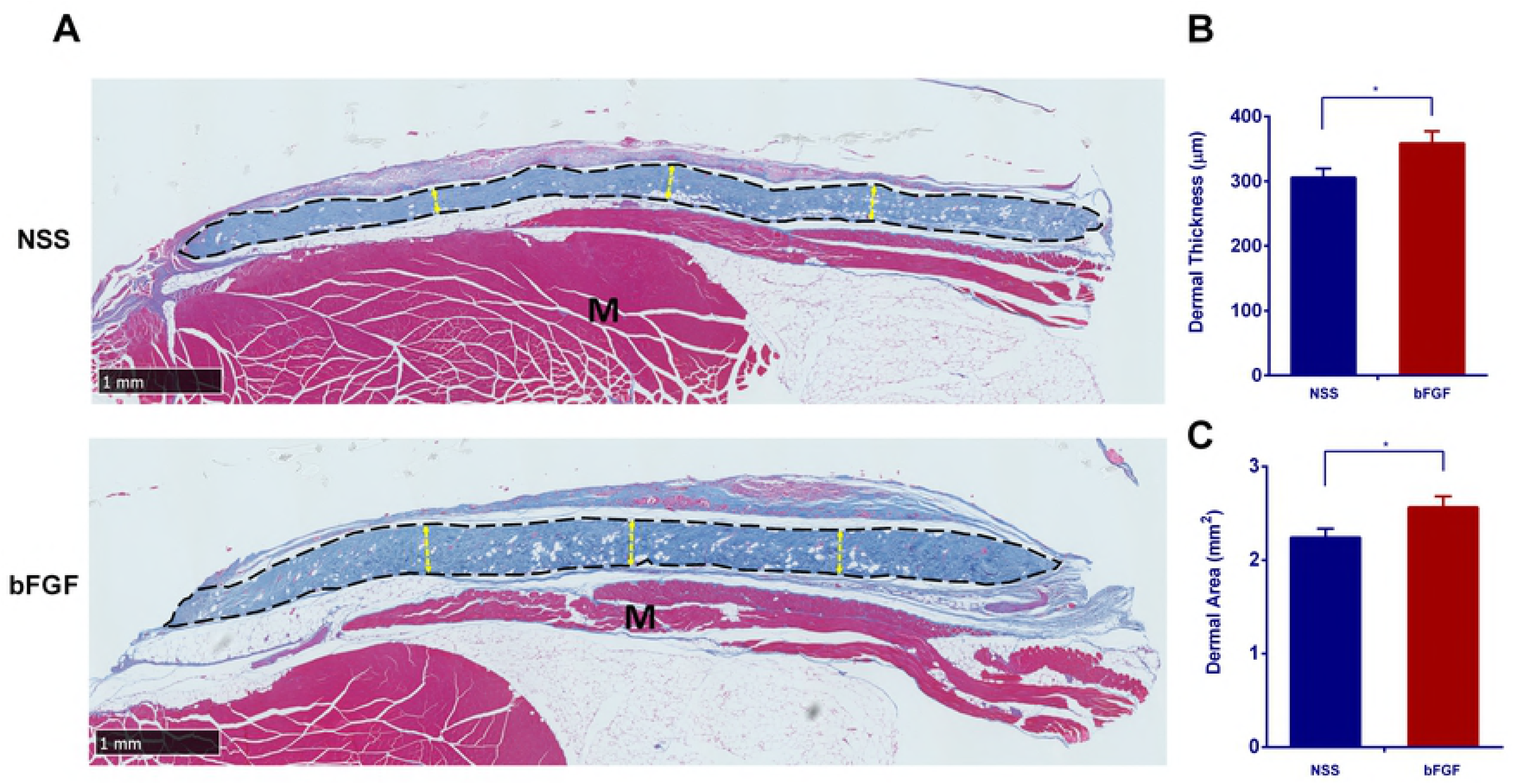
The assessment of dermal thickness and area after 8 weeks implantation. (A) A micrograph of the Azan staining at Week 8. The areas of the grafts are indicated by broken black lines. No signs of infection or necrosis were observed in either group. The yellow broken line indicates the evaluated thickness of the grafts, M: recipients’ muscle; Magnification: 2x, scale bar: 1 mm. (B) A comparison of the thickness of the grafts between the two groups at Week 8. The grafts in the bFGF group were significantly thicker than those in the NSS group (* p < 0.05). (C) A comparison of the area of the grafts between the two groups at Week 8. The dermis of the bFGF group was significantly larger than that in the NSS group (* p < 0.05).

## Discussion

GCMN are rare pigmented lesions most commonly defined as those > 20 cm in greatest dimension in adulthood or > 6-cm trunk lesion in an infant. Due to the increased risk of melanoma in such lesions, surgical excision in early childhood is recommended in order to prevent malignant transformation [32]. However, it has long been a challenge to completely excise the nevi, due to the limited availability of autologous skin.

In Japan, a CEA product (JACE^®^; Japan Tissue Engineering Co, Ltd., Gamagori, Japan) began to be applied for the treatment of GCMN in 2016; however, its take rate has not been satisfactory, mainly because the regeneration of the dermal component at the recipient site has not been established [33, 34]. To overcome this issue, we developed a novel treatment to inactivate the excised patient’s nevi by HHP and reconstruct the dermal component using the inactivated nevi and regenerate full-thickness skin in combination with autologous cultured epidermis [24]. The most important point of this method is to inactivate the nevus cells completely without damaging their native dermal structure, which supports sufficient strength and elasticity for the engraftment. We previously reported that HHP exceeding 200 MPa completely inactivated cells in the nevus tissue without damaging the structure of the ECM, while HHP exceeding 500 MPa damaged the epidermal basement membrane [25]. Although we found that the remaining cellular debris did not adversely affect the take of the inactivated skin [23, 24, 27, 35], the infiltration of fibroblasts from the wound bed into the inactivated skin took more than a week in a porcine autograft model, and the graft tended to be absorbed after four weeks [27, 35].

In the present study, we applied CGSs impregnated with bFGF to the inactivated skin in order to accelerate neovascularization to the inactivated skin and prevent the graft’s absorption after four weeks. bFGF has been well studied since the 1980s in order to clarify its pivotal role in wound regeneration through the acceleration of fibroblast proliferation and angiogenesis [1–3]. CGS is the scaffold that can sustain the release of bFGF for around 10 days *in vivo* [7, 12]. The optimal concentration of bFGF for impregnation to CGS was determined to be between 7 and 14 μg/cm^2^ [11, 36].

Among the many growth factors that contribute to the healing process, the most important positive regulators of angiogenesis are VEGF-A and FGF-2 (or bFGF) [37–39]. VEGF acts as an endothelial cell mitogen, chemotactic agent and inducer of vascular permeability, exhibiting unique effects on multiple components of the wound healing cascade, including angiogenesis and, recently, epithelization and collagen deposition [40]. Previous studies have suggested that bFGF may set the stage for angiogenesis during the first three days of wound repair, whereas VEGF may be critical for angiogenesis during granulation tissue formation from days 4 through 7 [40, 41]. Likewise, PDGF was the first growth factor shown to be chemotactic for cells migrating into the healing skin wound and has been suggested to be a major player in wound healing. In the present study, we evaluated the mRNA expressions of endogenous FGF-2, PDGF-A and VEGF-A in the inactivated grafts at Weeks 1 and 2. At Week 1, the mRNA level of FGF-2, PDGF-A and VEGF-A in the bFGF group was significantly higher than in the NSS group, indicating that the released bFGF from CGSs accelerated the cell infiltration from the wound bed to the inactivated skin and stimulated the expression of these proteins through the first week after grafting. Our CGSs are able to sustain the release of bFGF for 10 days, so such stimulation was no longer observed by Week 2. However, the infiltration of fibroblasts and capillaries into the inactivated skin was significantly accelerated at Week 2.

These findings suggest that the bFGF released from CGSs first stimulated the cells and then induced infiltration. At Week 8, we confirmed the successful take and survival of the inactivated skin histologically.

In the present study, we were unable to prepare murine cultured epidermis, so we could not confirm the survival of the cultured epidermis on the inactivated dermis. We will confirm this using inactivated human skin and human cultured epidermis *in vitro* or *in vivo* in the next step of our research.

## Conclusion

The survival of the inactivated skin by HHP at 200 MPa for 10 min was improved by the application of CGSs impregnated with bFGF. bFGF released from CGSs stimulated angiogenesis at the inactivated skin and improved the graft’s take at Week 8. The induction of angiogenesis and dermal regeneration in inactivated grafts through the sustained release of bFGF is suggested to be effective for improving the take rate of CEA application.

